# Chromosomal assembly and analyses of genome-wide recombination rates in the forest pathogenic fungus *Armillaria ostoyae*

**DOI:** 10.1101/794651

**Authors:** Renate Heinzelmann, Daniel Rigling, György Sipos, Martin Münsterkötter, Daniel Croll

## Abstract

Recombination shapes the evolutionary trajectory of populations and plays an important role in the faithful transmission of chromosomes during meiosis. Levels of sexual reproduction and recombination are important properties of host-pathogen interactions because the speed of antagonistic co-evolution depends on the ability of hosts and pathogens to generate genetic variation. However, our understanding of the importance of recombination is limited because large taxonomic groups remain poorly investigated. Here, we analyze recombination rate variation in the basidiomycete fungus *Armillaria ostoyae*, which is an aggressive pathogen on a broad range of conifers and other trees. We constructed a dense genetic map using 198 single basidiospore progeny from a cross. Progeny were genotyped at a genome-wide set of single nucleotide polymorphism (SNP) markers using double digest restriction site associated DNA sequencing (ddRADseq). Based on a linkage map of on 11,700 SNPs spanning 1007.5 cM, we assembled genomic scaffolds into 11 putative chromosomes of a total genome size of 56.6 Mb. We identified 1984 crossover events among all progeny and found that recombination rates were highly variable along chromosomes. Recombination hotspots tended to be in regions close to the telomeres and were more gene-poor than the genomic background. Genes in proximity to recombination hotspots were encoding on average shorter proteins and were enriched for pectin degrading enzymes. Our analyses enable more powerful population and genome-scale studies of a major tree pathogen.

## Introduction

Recombination shapes the evolution of chromosomes and the evolutionary trajectory of populations (Haenel *et al*, 2018; Otto and Lenormand, 2002). Crossovers enable the pairing and proper disjunction of homologous chromosomes during meiosis and are essential for the long-term maintenance of chromosomal integrity (Fledel-Alon *et al*, 2009; Hassold and Hunt, 2001). Loss of recombination on chromosomes is often associated with degenerative sequence evolution including gene loss and deleterious rearrangements. For example, the consequences of recombination cessation largely shaped the evolution of sex chromosomes and mating-type regions in animal, plants and fungi (Charlesworth *et al*, 2000; Wilson and Makova, 2009). Recombination also has a fundamental impact on the organization of genetic variation within populations. Recombination breaks up linkage between alleles at different loci, thereby generating novel combinations across loci that can be exposed to selection. Decreased linkage between loci increases the efficacy of selection and, hence, promotes adaptation (Hill and Robertson, 2009; Otto and Barton, 1997; Otto and Lenormand, 2002). However, recombination can also break up linkage between co-adapted alleles across loci, thereby creating a potential evolutionary conflict.

The study of the role of sex and levels of recombination is particularly important for our understanding of coevolutionary arms races in host-pathogen interactions. Host populations are thought to be under strong selection to maintain sexual reproduction to escape co-evolving pathogens by generating novel genotypes (Hamilton, 1980; Lively, 2010; Morran *et al*, 2011). Similarly, pathogens are under strong selection pressure to adapt to resistant hosts. In addition to mutation rates, the level of recombination is likely under selection in pathogen populations (Croll *et al*, 2015; Möller and Stukenbrock, 2017; Sánchez-Vallet *et al*, 2018). Notable cases of pathogen emergence driven by recombination include epidemic influenza viruses (Nelson and Holmes, 2007), typhoid fever caused by *Salmonella enterica* (Didelot *et al*, 2007; Holt *et al*, 2008) and toxoplasmosis caused by outcrossing *Toxoplasma gondii* strains (Wendte *et al*, 2010). Sexual reproduction is also prevalent in many fungal plant pathogens playing an important role in adaptive evolution (Möller and Stukenbrock, 2017). In particular in crop pathogens, the level of recombination was proposed as a predictor for the speed at which the pathogen will overcome host resistance (McDonald and Linde, 2002; Stukenbrock and McDonald, 2008). While pathogens of crop received significant attention to elucidate the organization of genetic variation and the impact of recombination on genome evolution (Croll *et al*, 2015; Stukenbrock and Dutheil, 2018), the role of recombination in the evolution of tree pathogens or saprophytes is still largely unknown.

An important group of fungal tree pathogens and saprophytes is represented by the basidiomycete genus *Armillaria*. The numerous fungi of this genus play an important role in the dynamics of forest ecosystems worldwide (Heinzelmann *et al*, 2019; Shaw III and Kile, 1991). With their ability to degrade all structural components of dead wood causing a white-rot, *Armillaria* species contribute significantly to nutrient cycling in forest ecosystems (Hood *et al*, 1991). Moreover, *Armillaria* species act as facultative pathogens infecting the root systems of healthy or weakened trees, and eventually cause tree mortality (Guillaumin *et al*, 2005). In timber plantations, the presence of Armillaria root disease causes substantial economic losses (Laflamme and Guillaumin, 2005), whereas in natural forest ecosystems the disease impacts forest succession, structure and composition (Bendel *et al*, 2006; Hood *et al*, 1991; McLaughlin, 2001). In the Northern Hemisphere, *Armillaria ostoyae* is of special importance. It is widely distributed in North America and Eurasia and recognized as an aggressive pathogen on a broad range of conifers and other trees (Anderson and Ullrich, 1979; Guillaumin *et al*, 1993; Morrison *et al*, 1985; Ota *et al*, 1998; Qin *et al*, 2007). *A. ostoyae* challenges current containment strategies and the search for new control strategies is ongoing (Heinzelmann *et al*, 2019).

*Armillaria* spp. have relatively large and recently expanded genomes (Aylward *et al*, 2017; Sipos *et al*, 2017). Recently, the genomes of a European and a North American *A. ostoyae* strain were published (Sipos *et al*, 2017). The genome assembly of the European strain (SBI C18/9) is of 60.1 Mb and split into 106 scaffolds. The genome assembly for the North American strain (28-4) is similar in length (58.0 Mb) but considerably more fragmented. However, none of the to date published *Armillaria* genomes is yet assembled to chromosome-scale sequences (Collins *et al*, 2013; Sipos *et al*, 2017; Wingfield *et al*, 2016). Expanded gene families in *Armillaria* include pathogenicity-related genes, enzymes involved in lignocellulose-degradation and *Armillaria*-specific genes with mostly unknown functions (Sipos *et al*, 2017). Interestingly, in comparison with other white-rot fungi, *Armillaria* shows an under-representation of ligninolytic gene families and an overrepresentation of pectinolytic gene families (Sipos *et al*, 2017). *A. ostoyae* is out-crossing and progeny populations were successfully used to identify the genetic basis of a major colony morphology mutant phenotype (Heinzelmann *et al*, 2017). However, further insights into genome evolution of *Armillaria* and the genetic basis of phenotypic traits are hampered by a lack of a dense recombination map and a fully finished reference genome.

In this study, we first aimed to establish a chromosome-scale assembly for *A. ostoyae* using a dense recombination map. Second, we aimed to test for variation in recombination rates within and among chromosomes to identify putative recombination hotspots. Finally, we analyzed genomic correlates of recombination rate variation including GC-content, gene density and content of transposable elements.

## Material and Methods

### Mapping population, construction of genetic map and comparison with reference genome

The mapping population used in this study consisted of 198 single basidiospore progeny of the diploid *A. ostoyae* strain C15 (WSL Phytopathology culture collection number: M4408). This strain was collected from a Scots pine (*Pinus sylvestris*) situated in a forest stand in the Swiss Plateau (Prospero *et al*, 2004). The haploid progeny were obtained from a single basidiocarp obtained *in vitro* as described previously (Heinzelmann *et al*, 2017). The haploid progeny and the diploid parent were genotyped at a genome-wide set of single nucleotide polymorphism (SNP) markers making use of double digest restriction site associated DNA sequencing (ddRADseq). The genetic map was constructed *de novo* using R/ASMap version 0.4-4 (Taylor and Butler, 2017) which is based on the MSTmap algorithm of Wu *et al* (2008). The significance threshold was set to *P =* 10^−5^. The marker order in the final genetic map was compared to the order in the genome of the haploid *A. ostoyae* strain SBI C18/9 (assembly version 2, May 2016, Sipos *et al*, 2017). Strain SBI C18/9 (WSL Phytopathology culture collection number: M9390) originates from Switzerland but is unrelated to strain C15.

### Construction of chromosome-scale sequences

The scaffolds of the reference genome were assembled into near chromosome-scale sequences, hereafter termed pseudochromosomes, based on the order of scaffolds within linkage groups. Scaffolds which were split by the genetic map into fragments mapping to different linkage groups or well separated regions (i.e. > 650 kb apart) of a linkage group were broken up into fragments. Emerging, unanchored sequences were removed. Scaffolds (or fragments thereof) that were joined into pseudochromosomes were separated by gaps of 10 kb. Scaffolds and scaffold fragments which were not oriented by the genetic map were orientated randomly. The completeness of both the original genome assembly and the pseudochromosomes and the corresponding gene annotations was compared with BUSCO version 3.1.0 (Simão *et al*, 2015) using the Basidiomycota dataset (library basidiomycota_odb9).

### Count and distribution of crossover events

The number of crossover events per progeny and pseudochromosome was extracted from the genetic map using the countXO function of the R/qtl package, version 1.40-8 (Broman *et al*, 2003). We used locateXO (R/qtl) to identify the position of crossover events and extract flanking markers. For each crossover event, we calculated the physical distance of the two flanking markers. To check for the presence of potential non-crossover (= non-reciprocal recombination events), the distance of two consecutive crossover events on a pseudochromosome was calculated. We assessed the minimal distance of crossover events as the physical distance between the first marker following a crossover and the last marker before the next crossover.

### Recombination rate variation along pseudochromosomes

For each pseudochromosome, the recombination rate was estimated in non-overlapping 20 kb segments as follows. First, genetic positions were linearly interpolated every 20 kb based on genetic and physical positions of markers using the approx function of the of the R package ‘stats’, version 3.4.0 (R Development Core Team, 2017). Next, the genetic distance per segment was calculated as the difference in genetic distance of the end and start point of the segment. Finally, the recombination rate per segment was obtained by dividing the interpolated genetic distance by the segment size. A segment size of 20 kb was considered appropriate because the physical distance between consecutive markers (excluding marker pairs with a distance of ≤ 400 bp to avoid spurious marker resolution through markers associated with the same restriction site) was less than 10 kb for ~50% of marker pairs, and less than 20 kb for ~75% of the marker pairs (Figure 1). We tested for heterogeneity of recombination rate along the pseudochromosomes by comparing the observed distribution of recombination rates per segment with the expected distribution using Fisher’s exact test. For this, the 20 kb segments were binned into categories of 0, 1, 2, 3, 4, 5, 6-15 cM. A Poisson distribution with lambda equaling the average cM per segment was used as the expected distribution. *P*-values were estimated by Monte Carlo simulations with 10^6^ replicates. This test was conducted for each pseudochromosome independently and all pseudochromosomes together.

**Figure 1.**
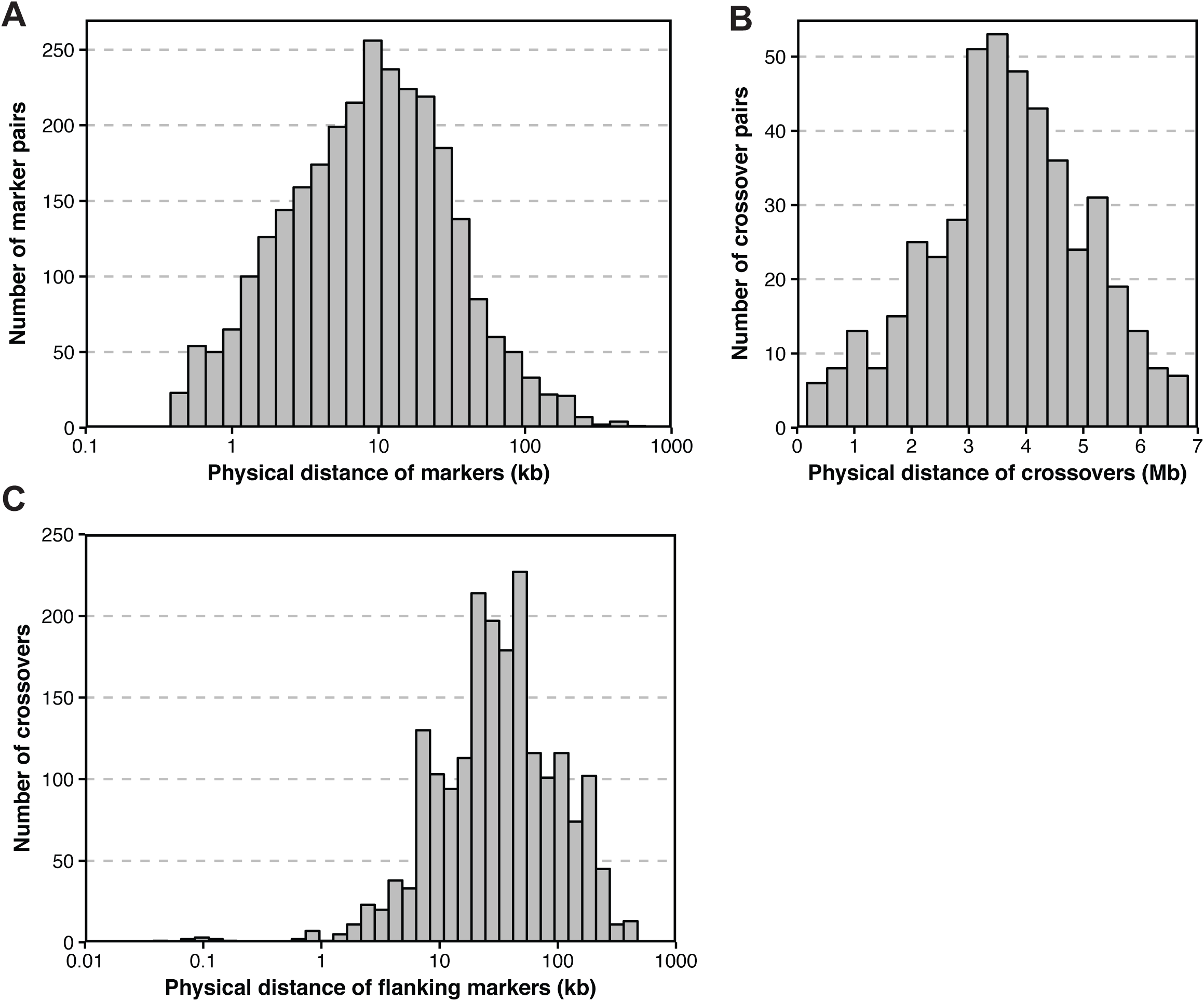
Resolution of the genetic map constructed for *Armillaria ostoyae* strain C15, distance of crossover events and accuracy of crossover placement. **A)** Physical distance between consecutive markers (marker pairs associated with the same restriction site excluded, see Materials and Methods) on pseudochromosomes LG 1 to LG 11. **B)** Physical distance of consecutive crossover events. The shortest distance recorded is 0.17 Mb. Because a major part of the left chromosome arm of pseudochromosome LG 11 may be missing, this pseudochromosome was excluded from this analysis. **C)** Physical distance of the two markers flanking a crossover. All pseudochromosomes were included.

### Identification and characterization of recombination hotspots

We identified recombination hotspots in the genome by searching for 20 kb segments with unusually high recombination rates (i.e. ≥ 200 cM/Mb). To account for the uncertainty in the identification of exact crossover locations, 20 kb segments with recombination rates ≥ 200 cM/Mb were conservatively extended by 15 kb on each side to define 50 kb recombination hotspot windows. In cases where two adjacent 20 kb segments had recombination rates ≥ 200 cM/Mb, one 50 kb hotspot centered on the two segments was created. Hotspots overlapping with assembly gaps were excluded. We assessed the correlation of GC-content, as well as gene and transposable element density with recombination hotspots. For this, the pseudochromosomes were divided into non-overlapping 50 kb segments. Segments were analyzed for GC-content and percentage of gene and transposable element coverage. Transposable elements were identified and annotated with RepeatModeler version 1.0.8 (A. F. A. Smit and R. Hubley, RepeatModeler Open-1.0 2008–2015; http://www.repeatmasker.org) and RepeatMasker version 4.0.5 (A. F. A. Smit, R. Hubley, and P. Green, RepeatMasker Open-4.0 2013– 2015; http://www.repeatmasker.org).

In addition, we assessed the correlation of recombination hotspots with certain gene properties and functions. Genes were functionally annotated using InterProScan version 5.19-58.0 (Jones *et al*, 2014). Protein families (PFAM) domain and gene ontology (GO) terms were assigned using hidden Markov models (HMM). Secretion signals, transmembrane, cytoplasmic, and extracellular domains were predicted using SignalP version 4.1 (Petersen *et al*, 2011), Phobius version 1.01 (Käll *et al*, 2004), and TMHMM version 2.0 (Krogh *et al*, 2001). A protein was conservatively considered as secreted only if SignalP and Phobius both predicted a secretion signal and no transmembrane domain was identified by either Phobius or TMHMM. Small secreted proteins were defined as secreted proteins shorter than 300 amino acids. Detailed gene annotations are provided in Supplementary Table S1. For plant cell wall degrading enzymes, i.e. enzymes involved in pectin, cellulose and hemicellulose and lignin degradation we relied on the annotations and categorization of Sipos *et al* (2017) (Supplementary Table S2). Similarly, we considered pathogenicity-related genes (including secondary metabolite genes) as identified by Sipos *et al* (2017) (Supplementary Table S3).

## Results

### Anchoring of the genome assembly to near chromosome-scale sequences

The genome of *A. ostoyae* strain C18/9 was sequenced using PacBio and Illumina sequencing technologies. PacBio reads were assembled into 106 scaffolds ranging from 5.0 kb to 6.4 Mb and polished using Illumina reads (Sipos *et al*, 2017). The total assembled genome size was 60.1 Mb. Here, we used a genetic map constructed for *A. ostoyae* strain C15 to assemble the genomic scaffolds into putative chromosomes (or pseudochromosomes). The genetic map was based on 11,700 high-quality SNP markers segregating in the mapping population. It contained 11 linkage groups and had a total length of 1007.5 cM (Heinzelmann *et al*, 2017). We were able to anchor 61 of the 109 scaffolds, which corresponds to 93% of the total sequence length of the genome assembly. The remaining 45 scaffolds were relatively short (5.0 to 338.3 kb). Overall, we observed a very high co-linearity of the marker order in the genetic map and the reference genome. Discrepancies were found in 13 scaffolds (scaffolds 1, 2, 4, 7, 9, 10, 12, 14, 16, 18, 27, 28 and 30). These scaffolds were split by the genetic map into 2 - 4 fragments that individually mapped either to different linkage groups or to well separated locations (i.e. > 650 kb apart) within the same linkage group (Supplementary Table S4). All scaffolds splits were supported by multiple markers from different restriction sites. In addition, we found that a part of scaffold 26 might be inverted or translocated in the genetic map relative to the reference genome.

Based on the genetic map, most of the anchored scaffolds (87.2%) could be oriented within pseudochromosomes (Table 1). Scaffolds (and fragments thereof) which could not be oriented (*n* = 10) were all short (44.1 to 246.7 kb). Each of the constructed pseudochromosomes was composed of 4 to 10 scaffolds or scaffold fragments. The total length of pseudochromosomes ranged from 3.3 to 7.0 Mb. The assembly into pseudochromosomes anchored 19 scaffolds with terminal telomeric repeats (TTAGGG)_≥7_, which were all located at the extremities of pseudochromosomes. An additional scaffold with terminal telomeric repeats could not be anchored. Overall, seven of the 11 pseudochromosomes had telomeres on both ends and the other four at one end, indicating that the pseudochromosomes represent in most cases nearly complete chromosomes (Table 1). The shortest pseudochromosome (LG 11) is substantially shorter than the others and might be missing a substantial portion of a chromosomal arm.

**Table 1.**
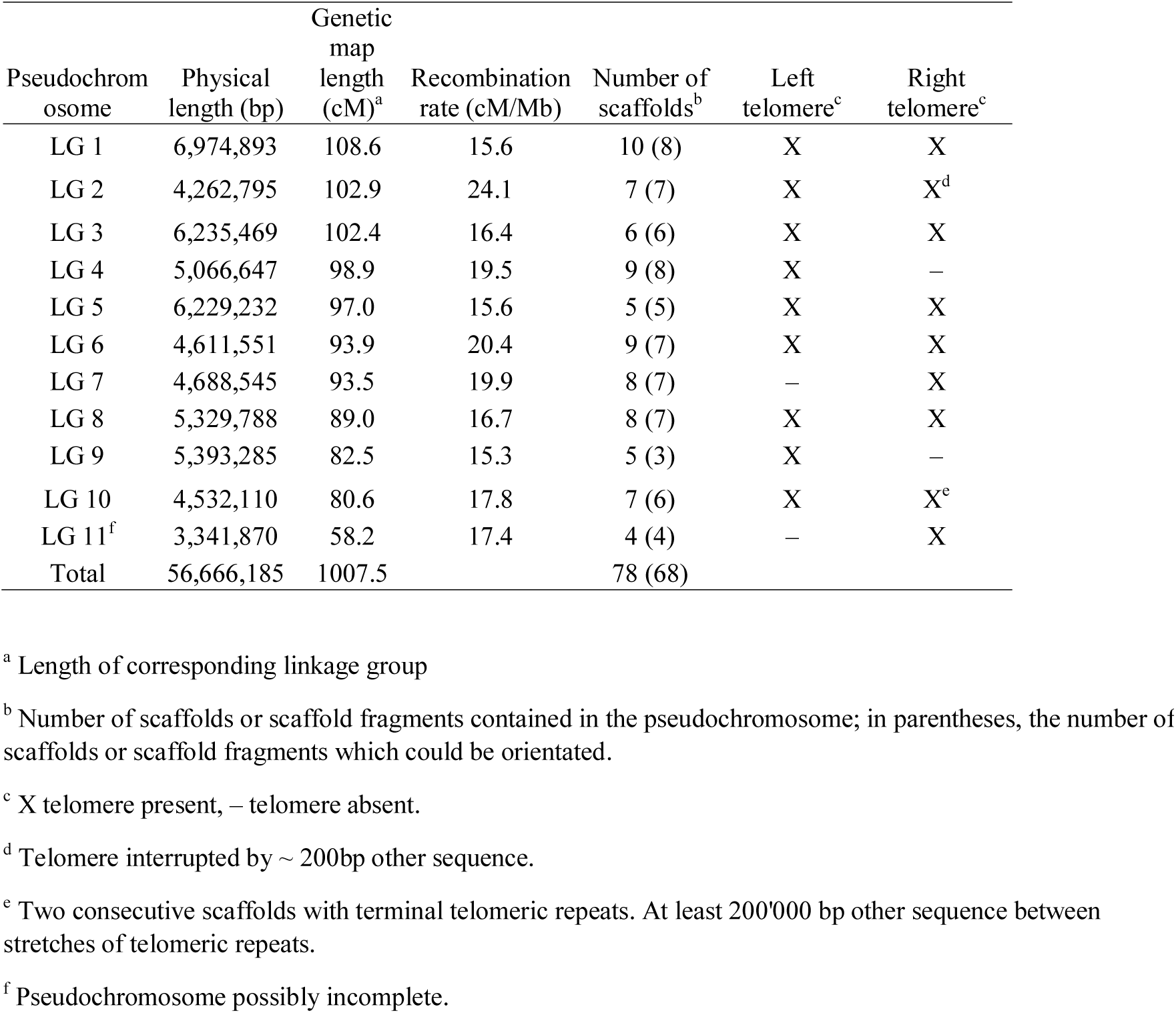
Overview of the pseudochromosomes constructed for *Armillaria ostoyae* based one the genetic map constructed in the progeny of the diploid *A. ostoyae* strain C15 and the genome assembly for the haploid *A. ostoyae* strain SBI C18/9.

### Frequency and distribution of crossover events

In total, we identified 1984 crossover events among all 198 progeny and 11 pseudochromosomes. The precision of crossover localization as determined by the spacing of SNP markers was below 20 kb for 33.6% and below 50 kb for 68.6% of crossover events (Figure 1). Consecutive crossover events on a chromosome were usually spaced far apart (Figure 1 and Supplementary Figure S1). On average, the distance between consecutive crossover events was at least 3.7 Mb with the closest two events being 0.17 Mb and the most distant 6.8 Mb apart. The large distance between crossover events indicates that most represent true crossovers, as non-crossovers are expected at much shorter distances. The possibly incomplete pseudochromosome LG 11 was discarded from the above analysis. The total number of crossover events observed per pseudochromosome varied from 115 (LG 11) to 214 (LG 1) (Table 2). The number of crossover events per progeny and chromosome varied from 0 to 3 with a median count of 1. Observing 3 crossovers on a chromosome was rare. We found no progeny with 3 crossovers on LG 2, LG 10 and LG 11 and a maximum of 7 progeny with 3 crossovers on LG 3. Pseudochromosome LG 11 had a very low mean crossover count compared to the other pseudochromosomes (0.58 *vs.* 0.80 - 1.08). On LG 11, only 4.5% of progeny were showing ≥ 2 crossover events compared to the other pseudochromosomes where 16 - 30% of progeny were showing 2 or 3 crossover events. This suggests that LG 11 is possibly missing a major part of a chromosomal arm without evidence what sequence constitutes the missing chromosomal fragment.

**Table 2.** Overview of crossover counts per linkage group in the progeny of the diploid *A. ostoyae* strain C15.

| Pseudochromosome | Occurrence of progeny with ... crossovers |  |  |  | Median crossover count | Mean crossover count | Crossover count per Mb | No. of crossovers total |
| --- | --- | --- | --- | --- | --- | --- | --- | --- |
|  | 0 | 1 | 2 | 3 |  |  |  |  |
| LG 1 | 47 | 91 | 57 | 3 | 1.00 | 1.08 | 0.15 | 214 |
| LG 2 | 44 | 106 | 48 | 0 | 1.00 | 1.02 | 0.24 | 202 |
| LG 3 | 52 | 98 | 41 | 7 | 1.00 | 1.02 | 0.16 | 201 |
| LG 4 | 51 | 101 | 45 | 1 | 1.00 | 0.98 | 0.19 | 194 |
| LG 5 | 56 | 92 | 44 | 6 | 1.00 | 1.00 | 0.16 | 198 |
| LG 6 | 60 | 97 | 37 | 4 | 1.00 | 0.92 | 0.20 | 183 |
| LG 7 | 57 | 101 | 39 | 1 | 1.00 | 0.92 | 0.20 | 182 |
| LG 8 | 60 | 102 | 35 | 1 | 1.00 | 0.88 | 0.17 | 175 |
| LG 9 | 70 | 96 | 30 | 2 | 1.00 | 0.82 | 0.15 | 162 |
| LG 10 | 74 | 90 | 34 | 0 | 1.00 | 0.80 | 0.18 | 158 |
| LG 11<sup>a</sup> | 92 | 97 | 9 | 0 | 1.00 | 0.58 | 0.17 | 115 |
| Total |  |  |  |  |  |  |  | 1984 |

<sup>a</sup> Pseudochromosome possibly incomplete.

### Heterogeneity of recombination rate along pseudochromosomes

The recombination rates estimated in non-overlapping 20 kb segments along pseudochromosomes were highly heterogeneous and varied from 0 to 737 cM/Mb (Figure 2). The median recombination rate was 2.5 cM/Mb. We tested whether the degree of heterogeneity along pseudochromosomes was deviating from a random distribution. When all pseudochromosomes (except pseudochromosome LG 11) were tested together, the recombination rate distribution was significantly different than a random distribution (Fisher’s exact test, *P* < 10^−6^, lambda of simulated distribution = 0.29). When tested individually, the recombination rate heterogeneity was significantly different than random on all but three pseudochromosomes (Table 3). In general, the highest recombination rates were observed towards the telomeres (Figure 2). We observed an inverse relationship of pseudochromosome length and recombination rate (r_Pearson_ = −0.77, *P* = 0.009, pseudochromosome LG 11 excluded).

**Table 3.** Recombination rate heterogeneity tests for individual pseudochromosomes. The observed distribution of recombination was compared to a Poisson distribution using Fisher’s exact test.

| Pseudochromosome | P-value<sup>a</sup> |  | Lambda<sup>b</sup> |
| --- | --- | --- | --- |
| LG 1 | < 10<sup>-3</sup> | *** | 0.25 |
| LG 2 | 0.534 | ns | 0.42 |
| LG 3 | < 10<sup>-3</sup> | *** | 0.27 |
| LG 4 | 0.006 | ** | 0.28 |
| LG 5 | 0.085 | ns | 0.29 |
| LG 6 | 0.031 | * | 0.34 |
| LG 7 | 0.033 | * | 0.30 |
| LG 8 | 0.002 | ** | 0.26 |
| LG 9 | 0.028 | * | 0.27 |
| LG 10 | 0.648 | ns | 0.27 |
| LG 11<sup>3)</sup> | - | - | - |

<sup>a</sup> \* $P < 0.05$ , \*\* $P < 0.01$ , \*\*\* $P < 0.001$ , ns: not significant.
<sup>b</sup> Lambda used for simulation of reference Poisson distribution. For details see Materials and Methods.
<sup>c</sup> No test was conducted as pseudochromosome is possibly incomplete.

**Figure 2.**
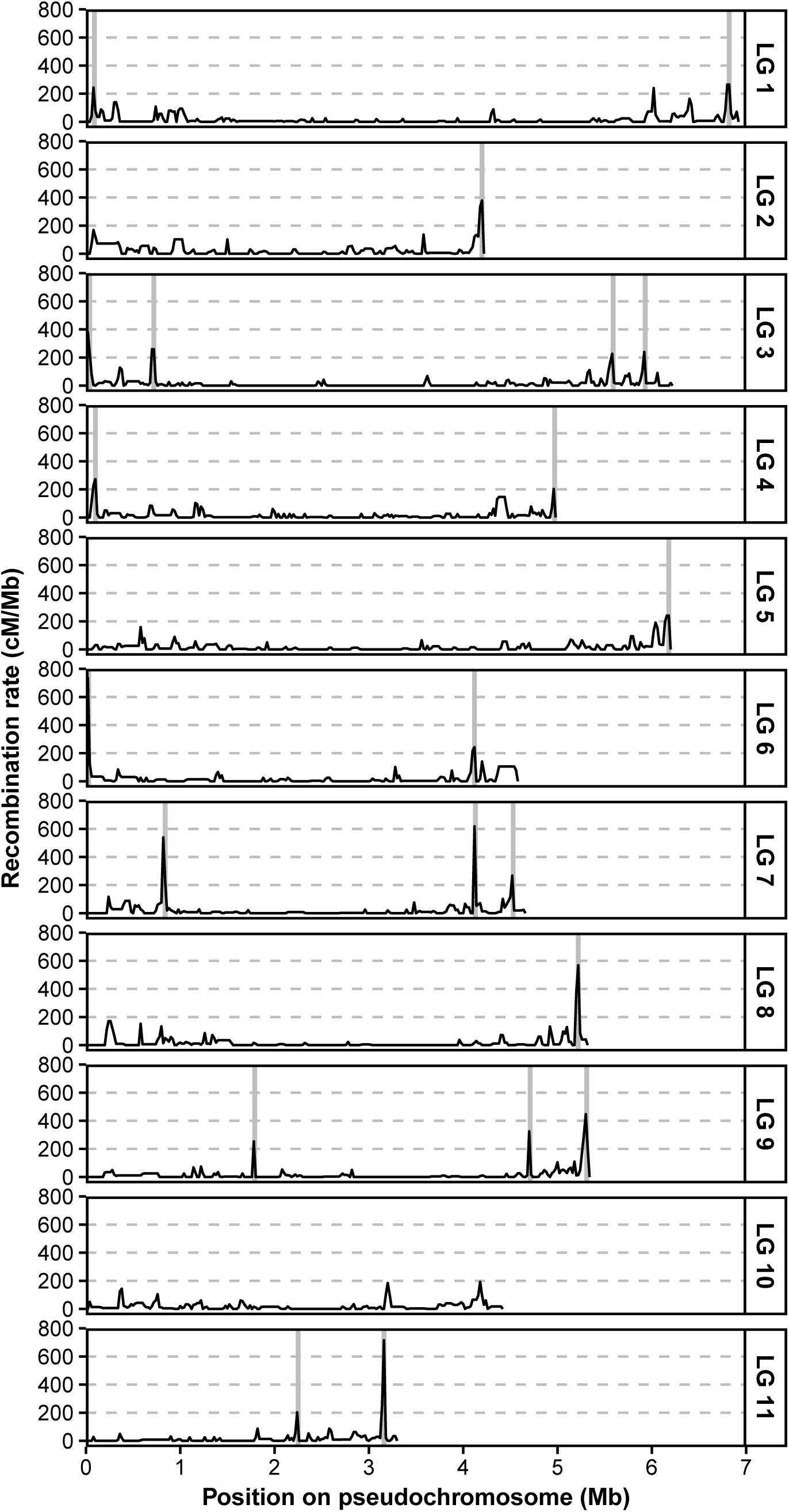
Recombination landscape of *Armillaria ostoyae* strain C15. Recombination rates were estimated in non-overlapping 20 kb segments. Vertical grey bars indicate the location of recombination hotspots defined as 50 kb windows centered on one or two adjacent 20 kb segments with recombination rates ≥ 200 cM/Mb. A potential recombination hotspot on LG 1 at 6 Mb was not considered a recombination hotspot because of an overlap with an assembly gap.

### Recombination hotspots

We defined recombination hotspots as narrow chromosomal tracts with the highest recombination rates. For this, we selected tracts of 20 kb chromosomal segments with recombination rates ≥ 200 cM/Mb. Both, the average and median recombination rate per 20 kb segment were with values of 17.6 cM/Mb and 2.5 cM/Mb, respectively, substantially lower. The probability to observe a 20 kb segment with a recombination rate of ≥ 200 cM/Mb by chance was *P* = 4.8 × 10^−4^ (Poisson distribution with lambda = 0.35 cM, which equals the average genetic distance per segment). In total, we identified 30 segments of 20 kb with recombination rates ≥ 200 cM/Mb. These segments represent only 1.1% of the analyzed genome sequence, but they accounted for 20.6% of the cumulative recombination rate. Overall, we found 19 distinct recombination hotspots on LG 1 to LG 10 (Figure 2). While pseudochromosome LG 11 was excluded from the above analyses, including LG 11 did not meaningfully affect the outcome of the above analyses (data not shown). On LG 11, we also identified two recombination hotspots (Figure 2).

### Association of recombination hotspots with sequence characteristics and gene content

*A. ostoyae* has a gene dense genome composed of 45.6% coding sequences (both when analyzing the complete scaffold assembly and the pseudochromosomes). The pseudochromosomes contained slightly less genes (21350 vs. 22705) compared to the complete scaffold assembly. The reduction in BUSCO completeness was reduced from 95.6% in the complete scaffold assembly to 95.2% in the pseudochromosomes. The content of transposable elements in the *A. ostoyae* genome is moderate (18.7% in the complete scaffold assembly and 14.5% in pseudochromosomes). Transposable elements tend to cluster and coincide with chromosomal regions with a lower GC-content and lower coding sequence density (Figure 3).

**Figure 3.**
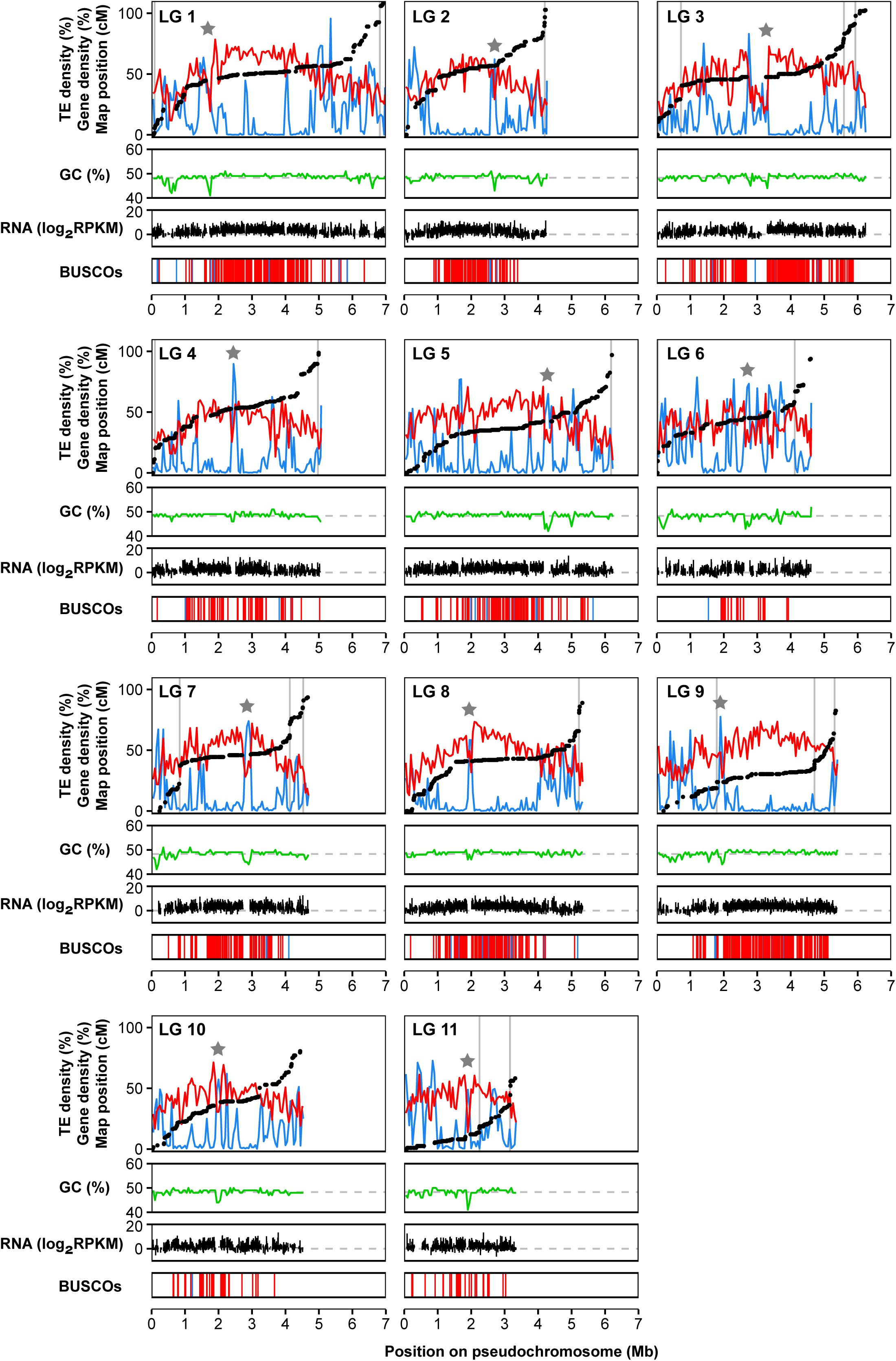
Characteristics of the *Armillaria ostoyae* pseudochromosomes. For each pseudochromosome, the first panel shows the genetic map position *vs.* the physical position of SNP markers (black dots). Gene density is shown in red and the density of transposable elements (TEs) is shown in blue. The position of recombination hotspots is indicated by grey vertical bars. Stars indicate approximate location of putative centromere regions. The second panel shows the GC content in green. The gray dashed line indicates the average GC content across all pseudochromosomes. Gene density, TE density and GC content were all estimated in non-overlapping 50 kb windows. The third panel shows gene expression levels in the cap of a fruiting body of the diploid *A. ostoyae* strain C18, which is the parental strain of the sequenced monosporous strain. Average gene expression among three biological replicates is shown. Expression data are retrieved from Sipos et al. (2017). RPKM = Reads per kilobase of transcript per million mapped reads. The fourth panel shows the position of BUSCO genes. Complete single copy BUSCO genes are shown in red and duplicated BUSCO genes are shown in blue.

We found that recombination hotspots (defined as 50 kb windows centered on identified hotspots) had a significantly lower density in coding sequences compared to the genomic background (32.4 ± 10.5% (± standard deviation) *vs.* 45.7 ± 13.4%; Mann–Whitney *U* test, *W* = 4482, *P* = 2.9 × 10^−5^) (Figure 4). The density of transposable elements in recombination hotspots was not significantly different to the genomic background (8.0 ± 8.1% *vs.* 14.2 ± 18.6%; Mann–Whitney *U* test, *W* = 9826, *P* = 0.798) (Figure 4). Transposable element densities varied widely among windows. The median hotspot window had a transposable element density of 4.8% compared to the 5.4% in the genomic background. GC-content was nearly identical in recombination hotspots and the genomic background (48.3 ± 0.8% *vs.* 48.4 ± 1.2%; Mann–Whitney *U* test, *W* = 8337, *P* = 0.145) (Figure 4).

**Figure 4.**
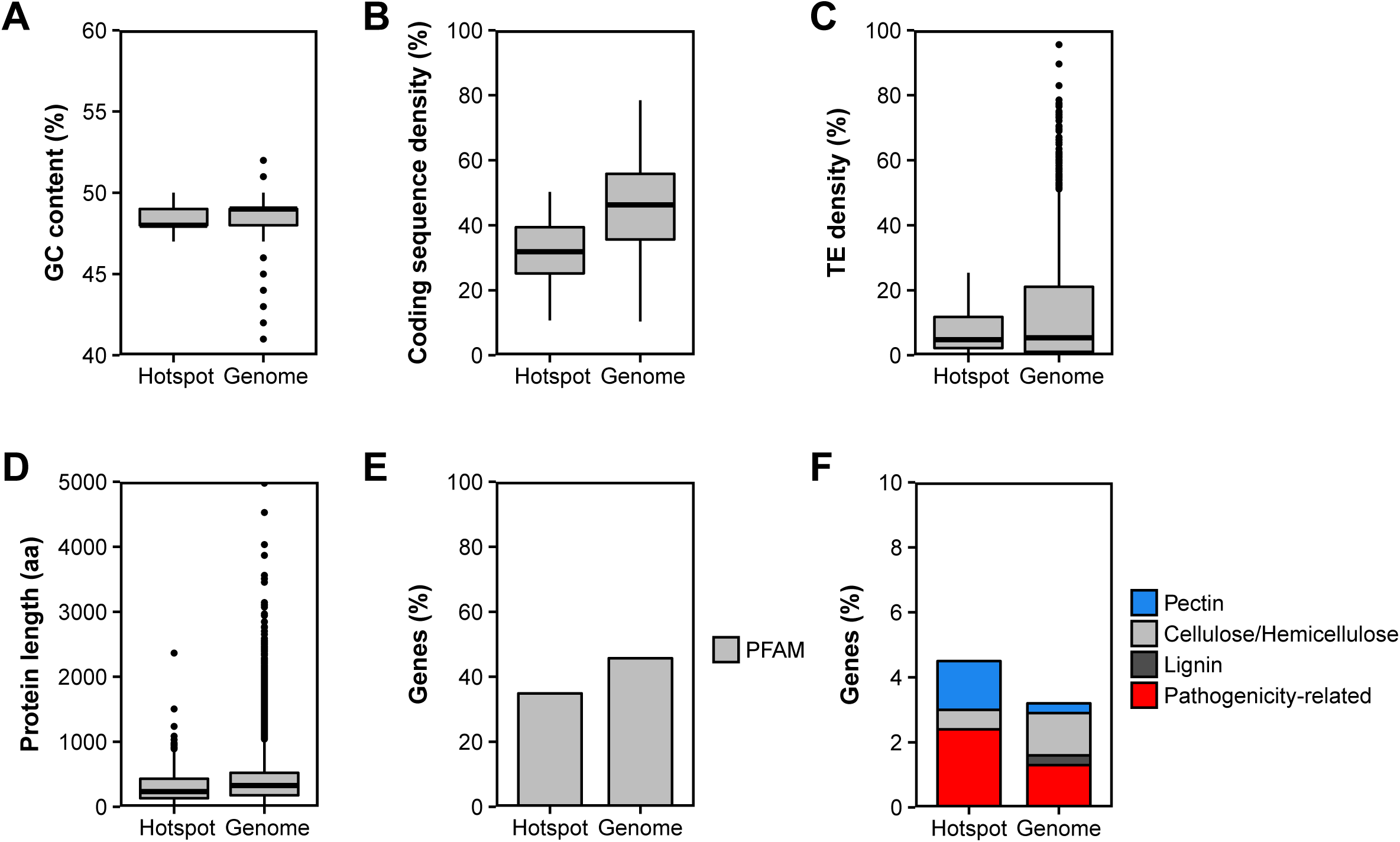
Characteristics of recombination hotspots in comparison to the genomic background. **A)** GC content, **B)** coding sequence density, **C)** density of transposable elements (TEs), **D)** protein length, **E)** percentage of genes with conserved domains (i.e. with PFAM annotation) and **F)** percentage of plant cell wall degrading genes (including pectin, cellulose, hemicellulose and lignin degrading genes) and pathogenicity-related genes. **A**-**C** were estimated for the genomic background in non-overlapping 50 kb windows.

We found that genes overlapping with recombination hotspots were encoding on average shorter proteins (Mann–Whitney *U* test, *W* = 2752400, *P* = 9.6 × 10^−8^) (Figure 4). Protein length averaged 322.8 ± 268.1 amino acids in recombination hotspots and 406.1 ± 339.4 amino acids in the genomic background. Proteins encoded in recombination hotspots were less likely to contain conserved PFAM domains (Fisher’s exact test, *P* = 8.2 × 10^−5^) compared to the genomic background (34.9% *vs.* 45.7%) (Figure 4). In the chromosomal context, we noted a lower density of genes with conserved PFAM domains at chromosome peripheries compared to chromosome centers (Figure 5). The frequency of genes encoding secreted proteins as well as small secreted proteins (< 300 aa) we found to be similar between recombination hotspots and the genomic background (Supplementary Table S5). Next, we analyzed plant cell wall degrading enzymes. Genes encoding pectin degrading enzymes were significantly overrepresented in recombination hotspots compared to the genomic background (Fisher’s exact test, *P* = 0.007) whereas genes encoding cellulose, hemicellulose and lignin degrading enzymes were similarly distributed among hotspots and the genomic background (Figure 4 and Supplementary Table S5). Pathogenicity-related genes (Sipos *et al*, 2017) tended to be more frequent in hotspots *vs.* non-hotspot regions, but the difference was not statistically significant (Supplementary Table S5). Overall, pathogenicity-related genes showed mostly a scattered distribution among all pseudochromosomes except for LG 6 where a large cluster of pathogenicity-related genes was observed (Figure 5). While pseudochromosome LG 11 was excluded from the above analyses, including LG 11 did not meaningfully affect the outcome of the above analyses (data not shown).

**Figure 5.**
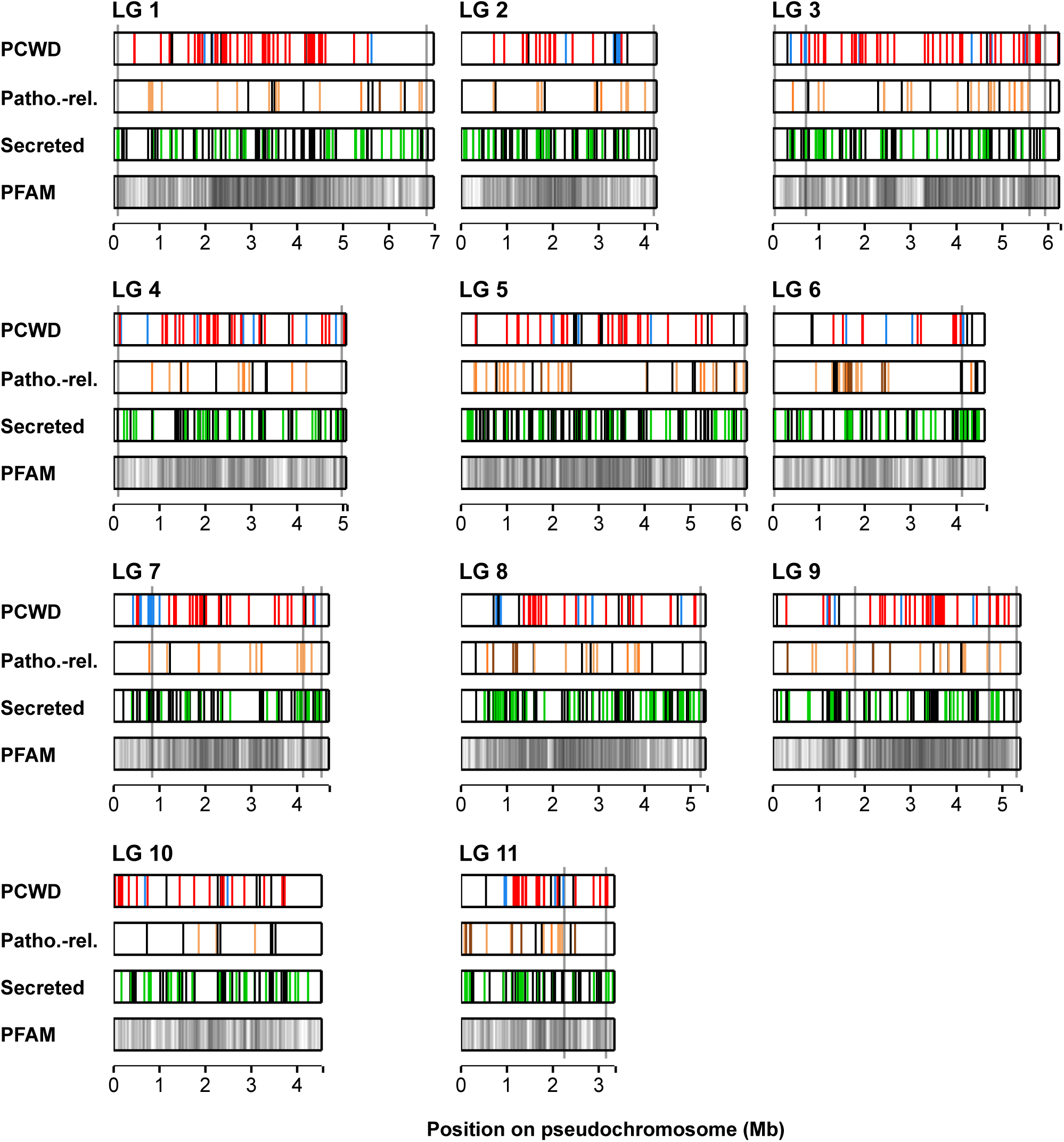
Distribution of genes encoding plant cell wall degrading (PCWD) enzymes (top bar), pathogenicity-related proteins (second bar), secreted proteins (third bar) and genes with conserved domains (i.e. PFAM annotation) (bottom bar) along the pseudochromosomes of *Armillaria ostoyae*. The following categories of plant cell wall degrading enzyme are distinguished: pectin degrading enzymes (blue), cellulose and hemicellulose degrading enzymes (red) and lignin degrading enzymes (black). Of the pathogenicity-related proteins, the three most frequent categories are highlighted: NRPS-like synthases (black), hydrophobins (dark brown) and carboxylesterases (medium brown). All other types of pathogenicity-related proteins are colored in light brown. Short secreted proteins (< 300 aa) are indicated in green, whereas all other secreted proteins are indicated in black. The density of genes with conserved domains is highest in dark areas and lowest in brighter areas. The locations of recombination hotspots are indicated by vertical gray bars spanning the horizontal bars.

## Discussion

We constructed a dense genetic map for *A. ostoyae* that enabled assembling a chromosome-scale reference genome. The presence of telomeric repeats on all but four pseudochromosomal ends indicates that nearly all chromosomes are completely assembled. Recombination rates increased from central regions towards the pseudochromosomal ends (*i.e.* telomeres). This further confirms the reliability of the chromosomal assembly. In addition, all chromosomes contain a putative centromere region of variable length (Figure 3), which is, as in other fungi, characterized by high transposable element density, low gene density, absence of gene transcription and reduced CG-content (Müller *et al*, 2019; Smith *et al*, 2012; Yadav *et al*, 2018). Putative centromere regions were located within chromosomal regions avoid of recombination, consistent with the findings from other fungi (Laurent *et al*, 2018; Mancera *et al*, 2008; Müller *et al*, 2019). The exact location and length of centromere regions, however, needs to be confirmed using chromatin immunoprecipitation sequencing (CHIPseq) as applied in other basidiomycetes (Yadav *et al*, 2018).

The previous assembly of the *A. ostoyae* genome into sub-chromosomal scaffolds was highly complete as assessed by BUSCO (Sipos *et al*, 2017). Even though we were unable to place ~ 7% of the total scaffold sequences, the unplaced scaffolds seem to contain mostly repetitive sequences. This was evident from the fact that our chromosome-scale assembly had only a very slightly reduced assembly completeness (95.2% *vs.* 95.6% of BUSCO genes). The transposable element content of our assembly is indeed quite lower compared to the scaffold-level assembly (14.5% *vs.* 18.7%). The status of the unplaced, repeat-rich scaffolds is difficult to assess. Our genetic map clearly lacked sufficient reliable markers to place small, repeat-rich scaffolds. We also identified a small number of discrepancies between the assembled scaffolds and the corresponding genetic map. These discrepancies were in all cases disjunctions of scaffolds and may be due to genetic differences between the sequenced strain (SBI C18/9) and the parental strain used for genetic mapping (C15). Some discrepancies may also stem from scaffold assembly errors. To fully resolve the causes for these discrepancies additional long-read sequencing is necessary. The pseudochromosome LG 11 is less complete and likely misses a substantial part of a chromosomal arm. This was evident from the short genetic map length and the markedly reduced number of progeny with at least 2 crossover events compared to the other pseudochromosomes (4.5% *vs.* 16 - 30%). The missing sequence may contain the rDNA cluster, which is challenging to assemble even with long-read sequencing and may constitute a substantial fraction of a fungal chromosome (Sonnenberg *et al*, 2016; Van Kan *et al*, 2017). The scaffold assembly of *A. ostoyae* contains a scaffold with three units of the rDNA repeat. However, we were unable to place this scaffold supporting the idea that our LG 11 assembly lacks the rDNA repeat. Alternatively, the missing chromosomal fragment may represent a major structural variation segregating between the strains SBI C18/9 and C15.

The identification of 11 pseudochromosomes (or linkage groups) provides the first estimate of the haploid chromosome number for an *Armillaria* species. Other species form the order Agaricales were found to have similar chromosome numbers: e.g. *Agaricus bisporus* (n = 13, Sonnenberg *et al*, 1996), *Coprinopsis cinerea* (n = 13, Muraguchi *et al*, 2003), *Pleurotus ostreatus* (n = 11, Larraya *et al*, 1999) or *Laccaria montana* (n = 9, Mueller *et al*, 1993). Given that our genetic map reached marker saturation and covers 93% of the scaffold-level assembly, the presence of additional chromosomes is highly unlikely. Karyotyping (*e.g.* by pulsed field gel electrophoresis) and high-density optical mapping would provide further confirmation of chromosome numbers and sizes, and likely resolve the placement of the remaining scaffolds. In particular, an optical map may help to resolve the size and position of the highly repetitive rDNA cluster (Van Kan *et al*, 2017).

The total size of the genetic map for *A. ostoyae* was 1007.5 cM and falls into the range of genetic map sizes observed for other basidiomycetes (Foulongne-Oriol, 2012). However, the total genetic map size depends on chromosome numbers and chromosomal recombination rates, which both vary substantially among fungal species. The *A. ostoyae* chromosomes all had a map length of 80.6-108.6 cM (with the exception of LG 11). This represents approximately two crossover events per bivalent and meiosis, which is consistent with the number of progeny observed with 0 (~25%), or 1 (~50%) or 2 (~25%) crossovers per chromosome. Chromosomal crossover counts vary considerably among fungal species. For example, in *A. bisporus* there is on average just one obligate crossover per bivalent for all chromosomes (Sonnenberg *et al*, 2016), whereas in *Saccharomyces cerevisiae* the average is ~6 crossovers per bivalent (Mancera et al, 2008). Interestingly, in some fungi there is a strong positive correlation between chromosomes size and the number of crossovers (Mancera *et al*, 2008; Roth *et al*, 2018), but we found no such apparent correlation in *A. ostoyae*.

The recombination landscape of *A. ostoyae* follows a canonical pattern, with increased recombination towards the peripheries of chromosomes and decreased recombination towards centromeres. The most striking deviations in these patterns are recombination hotspots. Such recombination hotspots are observed in many fungal species (Croll *et al*, 2015; Laurent *et al*, 2018; Müller *et al*, 2019; Roth *et al*, 2018; Van Kan *et al*, 2017), however their specific role in genome and gene evolution is still largely unknown. In the wheat pathogen *Zymoseptoria tritic* recombination hotspots may serve as ephemeral genome compartments favoring the emergence of fast-evolving virulence genes (Croll *et al*, 2015). Recombination hotspots in *A. ostoyae* were with two exceptions all located at the peripheries of chromosomes, where gene densities are low and gene functions are less conserved. From an evolutionary perspective, placing recombination hotspots distal from conserved housekeeping genes should be favorable given the mutagenic potential of hotspots. Interestingly, we found that genes involved in pectin degradation were enriched in recombination hotspots compared to the genomic background. Pectin is a major component of the plant cell wall and pectinolytic enzymes are among the first enzymes secreted by plant pathogens during host infection (Herbert *et al*, 2003). However, pectinolytic enzymes may also serve as effectors and induce plant defense reactions (Herbert *et al*, 2003). Hence, rapid evolution of pectinolytic enzymes may provide an advantage to *Armillaria* in its arms race with its hosts.

## Acknowledgments

Sequencing libraries were generated in collaboration with the Genetic Diversity Center (GDC) of ETH Zurich and sequenced by the Quantitative Genomics Facility of the Department of Biosystems Science and Engineering (D-BSSE) of ETH Zurich in Basel. The *A. ostoyae* genome project was funded by the European Union in the framework of the Széchenyi 2020 Program (GINOP-2.3.2-15-2016-00052) to GS and MM and by the WSL to GS.

## Data availability

Progeny sequencing data is available on the NCBI SRA under the BioProject accession PRJNA380873. The updated genome assembly was submitted to the European Nucleotide Archive (new accession number pending). The previous scaffold assembly can be retrieved from European Nucleotide Archive under the accession FUEG01000000.

## Supplementary information

Supplementary Figures and Tables are available.

## References

Anderson JB, Ullrich RC (1979). Biological species of *Armillaria mellea* in North America. Mycologia 71: 402–414.

Aylward J, Steenkamp ET, Dreyer LL, Roets F, Wingfield BD, Wingfield MJ (2017). A plant pathology perspective of fungal genome sequencing. IMA Fungus 8: 1–15.

Bendel M, Kienast F, Rigling D, Bugmann H (2006). Impact of root-rot pathogens on forest succession in unmanaged *Pinus mugo* stands in the Central Alps. Can J For Res 36: 2666–2674.

Broman KW, Wu H, Sen Ś, Churchill GA (2003). R/qtl: QTL mapping in experimental crosses. Bioinformatics 19: 889–890.

Charlesworth B, Harvey PH, Charlesworth B, Charlesworth D (2000). The degeneration of Y chromosomes. Philos Trans Royal Soc B 355: 1563–1572.

Collins C, Keane TM, Turner DJ, O’Keeffe G, Fitzpatrick DA, Doyle S (2013). Genomic and proteomic dissection of the ubiquitous plant pathogen, *Armillaria mellea*: toward a new infection model system. J Proteome Res 12: 2552–2570.

Croll D, Lendenmann MH, Stewart E, McDonald BA (2015). The impact of recombination hotspots on genome evolution of a fungal plant pathogen. Genetics 201: 1213–1228.

Didelot X, Achtman M, Parkhill J, Thomson NR, Falush D (2007). A bimodal pattern of relatedness between the *Salmonella* Paratyphi A and Typhi genomes: convergence or divergence by homologous recombination? Genome Res 17: 61–68.

Fledel-Alon A, Wilson DJ, Broman K, Wen X, Ober C, Coop G et al (2009). Broad-scale recombination patterns underlying proper disjunction in humans. PLOS Genet 5: e1000658.

Foulongne-Oriol M (2012). Genetic linkage mapping in fungi: current state, applications, and future trends. Appl Microbiol Biotechnol 95: 891–904.

Guillaumin JJ, Legrand P, Lung-Escarmant B, Botton B (eds) (2005). L’armillaire et le pourridié-agaric des végétaux ligneux. INRA: Paris, pp 487.

Guillaumin JJ, Mohammed C, Anselmi N, Courtecuisse R, Gregory SC, Holdenrieder O et al(1993). Geographical distribution and ecology of the *Armillaria* species in western Europe. Eur J Forest Pathol 23: 321–341.

Haenel Q, Laurentino TG, Roesti M, Berner D (2018). Meta-analysis of chromosome-scale crossover rate variation in eukaryotes and its significance to evolutionary genomics. Mol Ecol 27: 2477–2497.

Hamilton WD (1980). Sex versus non-sex versus parasite. Oikos 35: 282–290.

Hassold T, Hunt P (2001). To err (meiotically) is human: the genesis of human aneuploidy. Nat Rev Genet 2: 280–291.

Heinzelmann R, Croll D, Zoller S, Sipos G, Münsterkötter M, Güldener U et al(2017). High-density genetic mapping identifies the genetic basis of a natural colony morphology mutant in the root rot pathogen *Armillaria ostoyae*. Fungal Genet Biol 108: 44–54.

Heinzelmann R, Dutech C, Tsykun T, Labbé F, Soularue J-P, Prospero S (2019). Latest advances and future perspectives in *Armillaria* research. Can J Plant Pathol 41: 1–23.

Herbert C, Boudart G, Borel C, Jacquet C, Esquerre-Tugaye M, Dumas B (2003). Regulation and role of pectinases in phytopathogenic fungi. In: Voragen F, Schols H and Visser R (eds) Advances in pectin and pectinase research. Springer: Dordrecht, pp 201–220.

Hill WG, Robertson A (2009). The effect of linkage on limits to artificial selection. Genet Res 8: 269–294.

Holt KE, Parkhill J, Mazzoni CJ, Roumagnac P, Weill F-X, Goodhead I et al(2008). High-throughput sequencing provides insights into genome variation and evolution in *Salmonella typhi*. Nat Genet 40: 987.

Hood IA, Redfern DB, Kile GA (1991). *Armillaria* in planted hosts. In: Shaw III CG and Kile GA (eds) Armillaria root disease. Agricultural Handbook No. 691. USDA Forest Service: Washington D.C., pp 122–149.

Jones P, Binns D, Chang H-Y, Fraser M, Li W, McAnulla C et al(2014). InterProScan 5: genome-scale protein function classification. Bioinformatics 30: 1236–1240.

Käll L, Krogh A, Sonnhammer ELL (2004). A combined transmembrane topology and signal peptide prediction method. J Mol Biol 338: 1027–1036.

Krogh A, Larsson B, von Heijne G, Sonnhammer ELL (2001). Predicting transmembrane protein topology with a hidden markov model: application to complete genomes. J Mol Biol 305: 567–580.

Laflamme G, Guillaumin JJ (2005). L’armillaire, agent pathogène mondial: répartition et dégâts. In: Guillaumin JJ, Legrand P, Lung-Escarmant B and Botton B (eds) L’ armillaire et la pourridié-agaric des végétaux ligneux. INRA: Paris, pp 273–289.

Larraya LM, Perez G, Penas MM, Baars JJP, Mikosch TSP, Pisabarro AG et al(1999). Molecular karyotype of the white rot fungus *Pleurotus ostreatus*. Appl Environ Microbiol 65: 3413–3417.

Laurent B, Palaiokostas C, Spataro C, Moinard M, Zehraoui E, Houston RD et al(2018). High-resolution mapping of the recombination landscape of the phytopathogen *Fusarium graminearum* suggests two-speed genome evolution. Mol Plant Pathol 19: 341–354.

Lively CM (2010). A review of red queen models for the persistence of obligate sexual reproduction. J Hered 101: S13–S20.

Mancera E, Bourgon R, Brozzi A, Huber W, Steinmetz LM (2008). High-resolution mapping of meiotic crossovers and non-crossovers in yeast. Nature 454: 479–485.

McDonald BA, Linde C (2002). Pathogen population genetics, evolutionary potential, and durable resistance. Annu Rev Phytopathol 40: 349–379.

McLaughlin JA (2001). Impact of Armillaria root disease on succession in red pine plantations in southern Ontario. For Chron 77: 519–524.

Möller M, Stukenbrock EH (2017). Evolution and genome architecture in fungal plant pathogens. Nat Rev Micro 15: 756.

Morran LT, Schmidt OG, Gelarden IA, Parrish RC, Lively CM (2011). Running with the red queen: Host-parasite coevolution selects for biparental sex. Science 333: 216–218.

Morrison DJ, Chu D, Johnson ALS (1985). Species of *Armillaria* in British-Columbia. Can J Plant Pathol 7: 242–246.

Mueller GJ, Mueller GM, Shih L-H, Ammirati JF (1993). Cytological Studies in *Laccaria* (Agaricales). I. Meiosis and postmeiotic mitosis. Am J Bot 80: 316–321.

Müller MC, Praz CR, Sotiropoulos AG, Menardo F, Kunz L, Schudel S et al(2019). A chromosome-scale genome assembly reveals a highly dynamic effector repertoire of wheat powdery mildew. New Phytol 221: 2176–2189.

Muraguchi H, Ito Y, Kamada T, Yanagi SO (2003). A linkage map of the basidiomycete *Coprinus cinereus* based on random amplified polymorphic DNAs and restriction fragment length polymorphisms. Fungal Genet Biol 40: 93–102.

Nelson MI, Holmes EC (2007). The evolution of epidemic influenza. Nat Rev Genet 8: 196.

Ota Y, Matsushita N, Nagasawa E, Terashita T, Fukuda K, Suzuki K (1998). Biological species of *Armillaria* in Japan. Plant Dis 82: 537–543.

Otto SP, Barton NH (1997). The evolution of recombination: removing the limits to natural selection. Genetics 147: 879–906.

Otto SP, Lenormand T (2002). Resolving the paradox of sex and recombination. Nat Rev Genet 3: 252–261.

Petersen TN, Brunak S, von Heijne G, Nielsen H (2011). SignalP 4.0: discriminating signal peptides from transmembrane regions. Nat Methods 8: 785–786.

Prospero S, Holdenrieder O, Rigling D (2004). Comparison of the virulence of *Armillaria cepistipes* and *Armillaria ostoyae* on four Norway spruce provenances. Forest Pathol 34: 1–14.

Qin GF, Zhao J, Korhonen K (2007). A study on intersterility groups of *Armillaria* in China. Mycologia 99: 430–441.

R Development Core Team (2017). R: A language and environment for statistical computing. R Foundation for Statistical Computing. Vienna, Austria.

Roth C, Sun S, Billmyre RB, Heitman J, Magwene PM (2018). A high-resolution map of meiotic recombination in *Cryptococcus deneoformans* demonstrates decreased recombination in unisexual reproduction. Genetics 209: 567–578.

Sánchez-Vallet A, Fouché S, Fudal I, Hartmann FE, Soyer JL, Tellier A et al(2018). The genome biology of effector gene evolution in filamentous plant pathogens. Annu Rev Phytopathol 56: 21–40.

Shaw III CG, Kile GA (eds) (1991). Armillaria root disease. Agricultural Handbook No. 691. USDA Forest Service: Washington D.C., pp 233.

Simão FA, Waterhouse RM, Ioannidis P, Kriventseva EV, Zdobnov EM (2015). BUSCO: assessing genome assembly and annotation completeness with single-copy orthologs. Bioinformatics 31: 3210–3212.

Sipos G, Prasanna AN, Walter MC, O’Connor E, Bálint B, Krizsán K et al(2017). Genome expansion and lineage-specific genetic innovations in the forest pathogenic fungi *Armillaria*. Nat Ecol Evol 1: 1931–1941.

Smith KM, Galazka JM, Phatale PA, Connolly LR, Freitag M (2012). Centromeres of filamentous fungi. Chromosome Res 20: 635–656.

Sonnenberg ASM, de Groot PW, Schaap PJ, Baars JJP, Visser J, Van Griensven LJ (1996). Isolation of expressed sequence tags of *Agaricus bisporus* and their assignment to chromosomes. Appl Environ Microbiol 62: 4542–4547.

Sonnenberg ASM, Gao W, Lavrijssen B, Hendrickx P, Sedaghat-Tellgerd N, Foulongne-Oriol M et al(2016). A detailed analysis of the recombination landscape of the button mushroom *Agaricus bisporus* var. *bisporus*. Fungal Genet Biol 93: 35–45.

Stukenbrock EH, Dutheil JY (2018). Fine-scale recombination maps of fungal plant pathogens reveal dynamic recombination landscapes and intragenic hotspots. Genetics 208: 1209–1229.

Stukenbrock EH, McDonald BA (2008). The origins of plant pathogens in agro-ecosystems. Annu Rev Phytopathol 46: 75–100.

Taylor J, Butler D (2017). R Package ASMap: Efficient Genetic Linkage Map Construction and Diagnosis. J Stat Softw 79: 1–29.

Van Kan JAL, Stassen JHM, Mosbach A, Van Der Lee TAJ, Faino L, Farmer AD et al(2017). A gapless genome sequence of the fungus *Botrytis cinerea*. Mol Plant Pathol 18: 75–89.

Wendte JM, Miller MA, Lambourn DM, Magargal SL, Jessup DA, Grigg ME (2010). Self-mating in the definitive host potentiates clonal outbreaks of the apicomplexan parasites *Sarcocystis neurona* and *Toxoplasma gondii*. PLOS Genet 6: e1001261.

Wilson MA, Makova KD (2009). Genomic analyses of sex chromosome evolution. Annu Rev Genom Hum G 10: 333–354.

Wingfield BD, Ambler JM, Coetzee MPA, de Beer ZW, Duong TA, Joubert F et al(2016). Draft genome sequences of *Armillaria fuscipes*, *Ceratocystiopsis minuta*, *Ceratocystis adiposa*, *Endoconidiophora laricicola*, *E. polonica* and *Penicillium freii* DAOMC 242723. IMA Fungus 7: 217–227.

Wu YH, Bhat PR, Close TJ, Lonardi S (2008). Efficient and accurate construction of genetic linkage maps from the minimum spanning tree of a graph. PLOS Genet 4: e1000212.

Yadav V, Sun S, Billmyre RB, Thimmappa BC, Shea T, Lintner R et al(2018). RNAi is a critical determinant of centromere evolution in closely related fungi. Proc Natl Acad Sci USA 115: 3108–3113.

