## Supplementary Figure S1 for "Chromosomal assembly and analyses of genome-wide recombination rates in the forest pathogenic fungus *Armillaria ostoyae*"

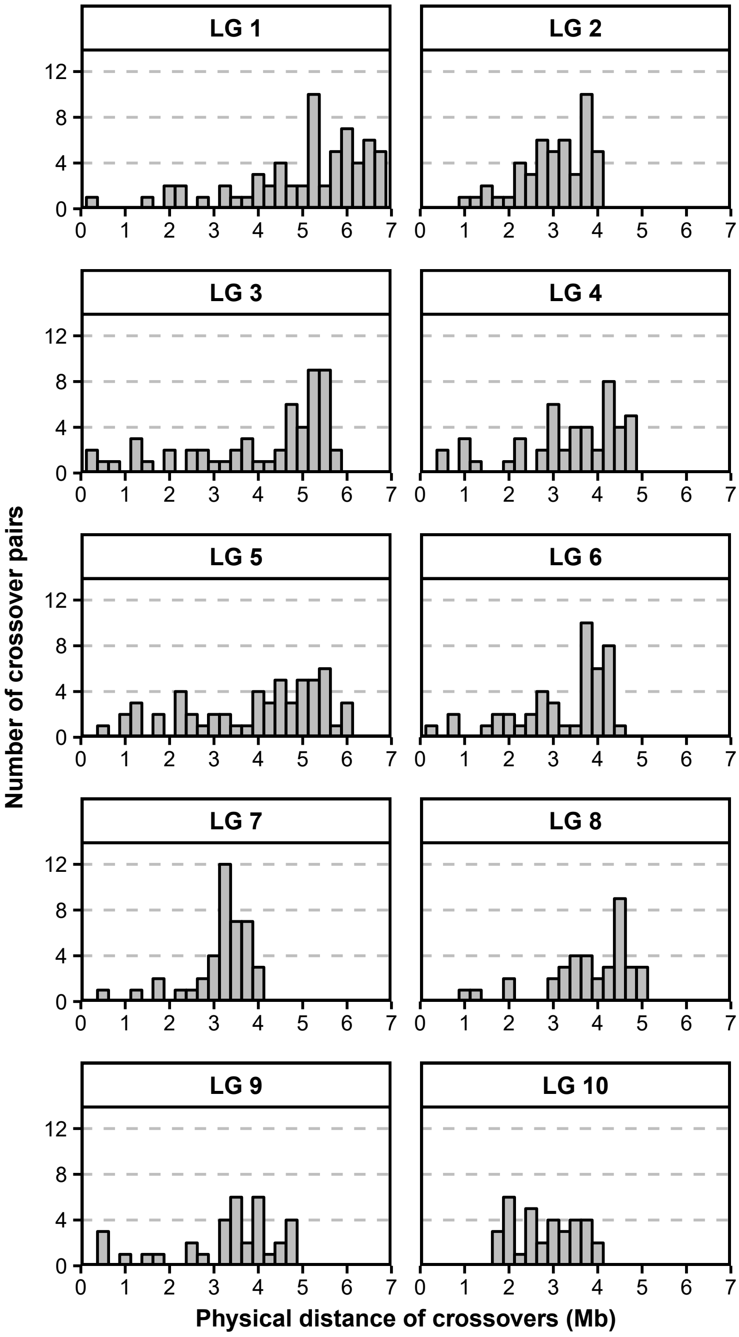


**Supplementary Figure S1.** Physical distance of consecutive crossover events on each of the pseudochromosomes across the 198 analyzed haploid progeny of *Armillaria ostoyae* strain C15. Because a major part of the left chromosome arm of pseudochromosome LG 11 is likely missing, no panel is shown for this pseudochromosome.
